# Near-equiprobable binary branching decisions underlie filament patterning in the moss *Physcomitrium patens*

**DOI:** 10.64898/2026.03.12.711315

**Authors:** Jeanne Abitbol-Spangaro, Bruno Chapuis, Christophe Godin, Yoan Coudert

**Author notes:** These authors contributed equally (Christophe Godin, Yoan Coudert).

## Abstract

Branching forms are ubiquitous in nature and have evolved repeatedly across scales and species. An important goal of developmental biology remains to identify similarities and differences in the regulatory mechanisms underlying branching. Here, we investigate the branching filaments that form upon spore germination in mosses, using *Physcomitrium patens* as a model species. To identify the macroscopic rules governing filament patterning, we developed a pipeline to acquire high-resolution 3D images of whole sporelings, reconstruct filament architecture at single-cell resolution, and formalize cell organization using mathematical tree representations. Our quantitative analysis reveals that branch patterning in moss filaments can be captured by a simple probabilistic model in which subapical cells have a near-equal probability of producing – or not producing – a side-branch between successive apical cell divisions. This framework provides a quantitative basis for comparing the developmental rules driving branching morphogenesis within and beyond the plant kingdom.

## Introduction

Branching is a fundamental developmental strategy that has emerged repeatedly during the evolution of biological species. In mobile organisms, branching generally occurs at the level of specific cells, such as neurons^1^, or organs, such as the vascular system^2^. In contrast, in sessile organisms, including corals, fungi, algae and plants, it manifests at the level of the whole organism, enabling them to optimize interactions with their local environment^3,4^. Its ubiquity across kingdoms and scales makes branching a striking example of convergent evolution. Identifying similarities and differences in the regulatory mechanisms underlying branching across species has therefore been an important focus of evolutionary and developmental biology. At the cellular and molecular levels, mechanisms driving branching morphogenesis can differ substantially. For instance, cell migration is a key factor in tissue remodeling and branching processes in animals^5,6^, whereas in plants, cells maintain fixed relative positions and development occurs primarily through cell growth and division. Yet, at the macroscopic scale, common regulatory mechanisms have been identified. For example, apical dominance, whereby the apex of a growing axis inhibits branch initiation at a distance, has been described in animals, such as colonial hydroids^7^, in fungi^8^, in algae^9^, as well as in the leafy shoots of land plants^10,11^.

During the diversification of plant body plans, the innovation of branching in filaments was likely a key step preceding the innovation of three-dimensional (3D) leafy shoots^12^. In mosses, the first tissue to develop following spore germination consists of branching filaments known as protonemata. These filaments initially develop as one-dimensional (1D) uniseriate cell files that rapidly branch, from which a small proportion gives rise to erect, 3D leafy shoots^13^. This developmental transition from 1D prostrate uniseriate filaments to 3D erect leafy shoots has sometimes been interpreted as recapitulating the innovation of 3D growth in the evolution of land plants^14^. For this reason, the filamentous stage of the moss life cycle may provide valuable insights into the evolution of branching morphogenesis in plants, as well as into the convergent evolution of branching in filamentous bodies across kingdoms^15^.

In the model moss *Physcomitrium patens*, filaments have traditionally been described by observing the periphery of well-developed (*i*.*e*., several week-old) gametophytes^16,17^. These filaments exhibit a stereotypical architecture marked by a gradient of cell identity. Caulonemal cells – elongated, with oblique division planes and relatively low chloroplast density – are localized at or near the apex, whereas chloronemal cells – relatively shorter, with perpendicular division planes and higher chloroplast density – are found closer to the base. Side-branches are typically distributed along a basitonic gradient^18^, with their length increasing along the apical-basal axis of the parent filament, conferring a characteristic “triangular” morphology^15,19^. Nevertheless, the developmental rules governing the establishment of this morphology remain largely unknown.

To date, protonemata have been characterized as tip-growing filaments, with growth occurring locally at the tip of apical cells, independently of cell identity^20^. Branching occurs through the establishment of a new growing tip that emerges laterally as a bulge in a subapical cell, generally on the distal side relative to the filament base^21^. However, beyond these local descriptions, an integrative view of branch patterning during protonemal development is still missing. Peripheral filaments of well-developed colonies, which are typically studied, are likely regulated by multiple concurrent biological processes integrated since the onset of gametophyte formation. Characterizing filament development at early stages therefore holds the potential to reveal the fundamental rules governing branching pattern establishment.

Using the moss *P. patens* as a model, this work aims to provide a quantitative description of branch patterning during the early stages of protonemal development, with the goal of identifying a minimal set of rules capturing its core morphogenetic principles.

## Results

### Light-sheet microscopy imaging reveals sporeling architecture at cellular resolution

To elucidate the macroscopic rules governing moss filament development, we developed a novel experimental pipeline to grow and image sporelings during their early developmental stages. This approach enabled detailed morphological and architectural analyses at cellular resolution. Spores from a transgenic line expressing a fluorescently tagged plasma membrane protein were germinated in liquid BCDAT medium (Figure 1A). After 4-7 days of growth, sporelings were imaged using light-sheet microscopy, allowing the acquisition of high-resolution 3D images capturing the entire filamentous structure (Figures 1B-C). We then used Arivis software to segment cells, which were individually labeled (Figure 1D). From these segmentations, we reconstructed the cellular network as a tree structure, in which nodes represent cells and edges represent cell division events (Figure 1E). Trees were rooted at the spore, and branching order was manually assigned based on visual inspection of the 3D images. Using this pipeline, we successfully segmented and reconstructed a total of 33 sporelings, each composed of 20 to 70 cells (Figures S1 and S2).

**Figure 1:**
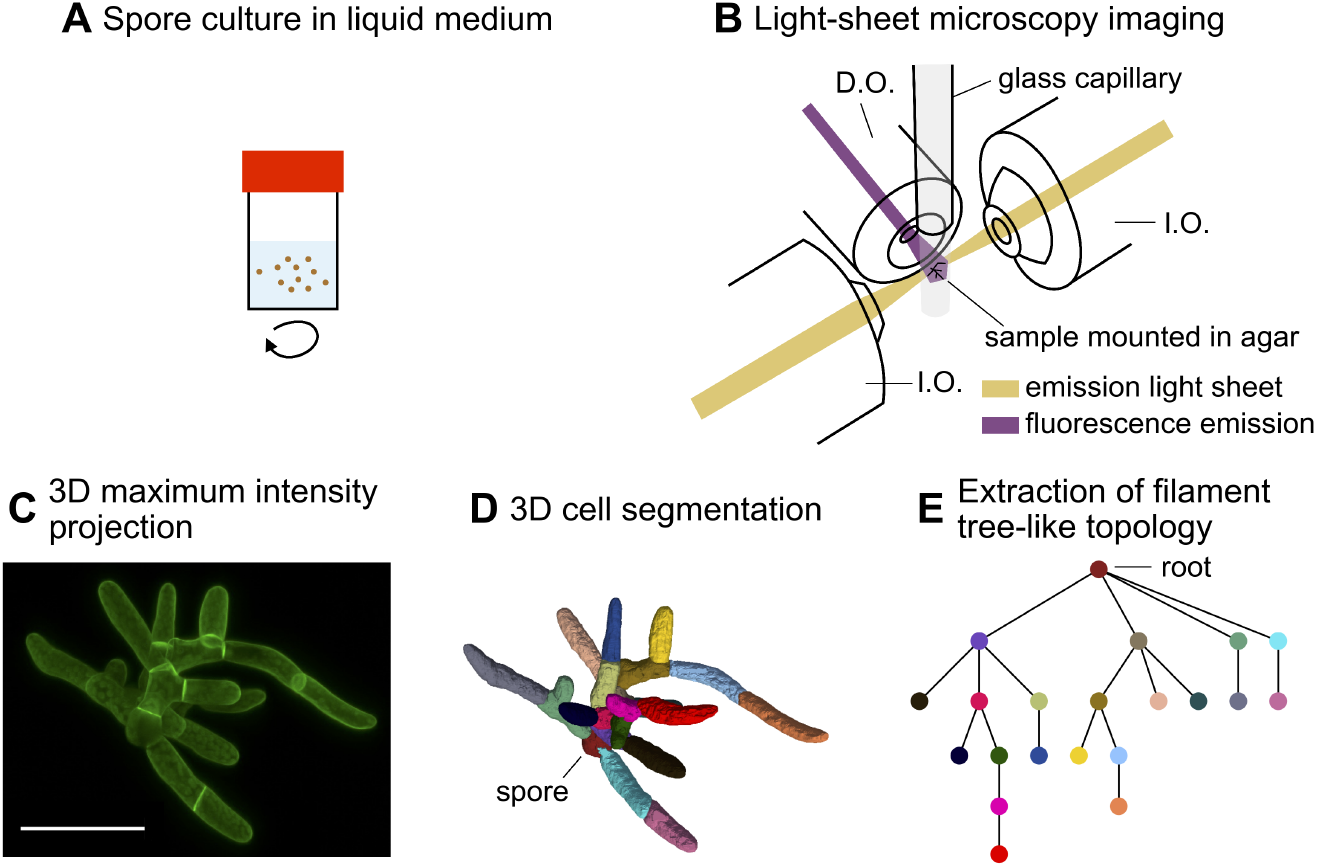
Data acquisition pipeline. **(A)** Schematic diagram of spore culture conditions, showing spores cultivated in liquid BCDAT medium under agitation. **(B)** Schematic diagram of the imaging set-up showing a single sporeling mounted in low-melting agarose inside a glass capillary for light-sheet microscopy. Light-sheet microscopy allows the acquisition of high-resolution 3D images using two Illumination Objectives (I.O.) that generate an emission light-sheet (yellow), and a Detection Objective (D.O.) that collects the fluorescence emission (purple). **(C)** 3D maximum intensity projection of the plasma membrane signal from a single sporeling. Scale bar = 100 *µ*m. **(D)** 3D cell segmentation of the sporeling shown in (C). **(E)** Tree-like representation of the sporeling architecture shown in (C, D). Nodes correspond to cells and edges correspond to cell division events. Node colors indicate cell labels consistent with panel (D). The root of the tree represents the spore, as indicated in (D).

### Early sporelings consist exclusively of chloronemal cells

To investigate cell identity in early sporelings, we first examined their overall morphology. Visually, most cells were relatively short and exhibited high chloroplast density, both characteristic features of chloronemal cells. We attempted to detect more subtle differences by quantifying parameters expected to distinguish chloronemal from caulonemal identity. Specifically, we plotted the distribution of cell length and chloroplast density along filaments with respect to both distance from the filament tip (measured in cell number) and branching order^19,22,23^ (Figures S3A-F). Pearson correlation analyses indicated that variability in cell length and chloroplast density could not be reliably explained by distance from the filament tip or branching order (Figures S3G-H, J). Unexpectedly, chloroplast density negatively correlated with distance from the filament tip (Figure S3I), whereas a positive correlation would be expected during the early chloronema-to-caulonema transition^20,23^, as reported previously. This observation suggests that chloroplast density gradually increases during early sporeling development, likely through regulatory mechanisms independent of the canonical chloronema-to-caulonema transition. Overall, these results indicate that all the analyzed filaments consisted exclusively of chloronemal cells, and the caulonemal transition had not yet occurred. The presence of a single cell type (*i*.*e*., chloronemata) in our dataset enabled branching pattern analyses to be performed consistently across sporelings.

### Filaments exhibit basitonic branching patterns

To quantify branching patterns, each cell file of the same branching order was treated as an individual filament across all sporelings, and side-branch number and length (measured in cell number) was plotted against cell position along the filament axis (Figures 2A-C). The first side-branch from the apex typically formed on the second sub-apical cell. At the population level (*i*.*e*., pooling all individual filaments), side-branch length increased approximately linearly (slope = 0.52), with branches closer to the spore being longer than those nearer the apex (Figure 2B). This basitonic branching pattern is consistent with sequential branch initiation followed by continuous growth. Additionally, the general absence of side-branches on the first sub-apical cell indicates that the competence for side-branch production is generally acquired only at the second sub-apical position, suggesting a developmental delay or inhibition. We observed that side-branch number also increased nearly linearly from the apex to the base (slope = 0.49), reaching up to four side-branches on the basalmost cells (Figure 2C). This reveals a higher branching competency than previously reported in the literature^13,16,19^, suggesting that the developmental stage of sporelings and the culture conditions are important determinants of cell branching potential.

**Figure 2:**
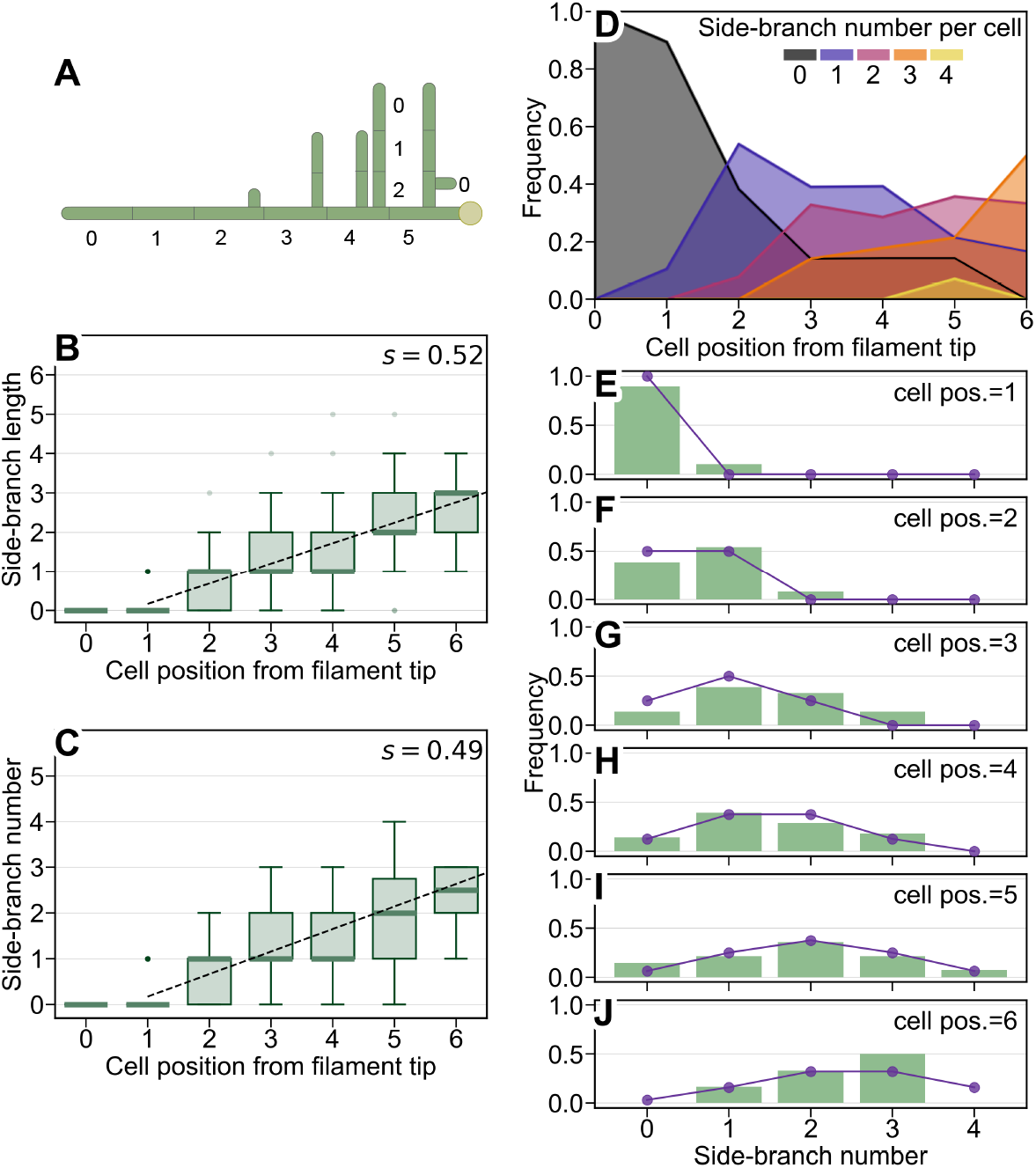
Spore-derived filaments display basitonic branching patterns captured by a binomial distribution. **(A)** Schematic diagram of a branched filament with cell numbering from filament tip. The spore is represented by a beige dot. **(B)** Box plots showing that side-branch length distribution as a function of cell position from the filament tip, as represented in (A), increases approximately linearly with slope *s* = 0.52. **(C)** Box plots showing that the number of side-branches per cell as a function of cell position from the filament tip, as represented in (A), increases approximately linearly with slope *s* = 0.49. **(D)** Intensity diagram of the number of side-branches per cell as a function of cell position from filament tip, as represented in (A). **(E-J)** Histograms of side-branch number distribution for cells at positions 1-6 (E to J, respectively) from the filament tip. Histograms are overlaid with expected frequencies from a binomial distribution with probability *p* = 0.5 and trial number *k* corresponding to cell position 1 (purple dots). *n*=405, 213, 116, 65, 28, 14, 6 cells for positions 0-6 from the filament tip.

### Filament branching is a stochastic process

To further characterize the mathematical properties underlying these branching patterns, we plotted the frequency of side-branch number per cell against cell position along individual filaments, represented either as an intensity diagram (Figure 2D) or as histograms for each cell position (Figures 2E-J). We noticed that the resulting distributions closely resembled those of a binomial distribution law, which describes the number of successes in a series of *k* independent trials with probability *p*, with cell position corresponding to the number of trials. Based on this observation, we hypothesized that branch initiation can be modeled as a sequence of draws with constant probability of success, with parent cell position corresponding to the number of trials, such that the second sub-apical cell has undergone one trial, the third sub-apical cell two trials, and so on. Under this hypothesis, the expectancy of side-branch number *E*[*X*] is given by the product of the success probability *p* and the number of trials *k* such that *E*[*X*] = *pk*, allowing *p* to be estimated from the slope of side-branch number as a function of trial number (*i*.*e*. parent cell position). The above results therefore indicate a branching probability *p* = 0.49 (Figure 2C). Consistently, we found that the observed frequency of side-branch number per cell could be approximated by a binomial distribution with *p* = 0.5 and *k* = cell position*−*1 (purple dots in Figures 2E-J). Together, these results suggest that filament cells have a near-equal chance of initiating or not initiating a side-branch following each apical cell division. This implies that branching during early sporeling development behaves as a stochastic process consistent with a binomial model, in which each cell along a filament axis – beyond the first sub-apical cell – may produce a branch in a cell-autonomous manner.

### A probabilistic model of filament growth and branching captures average distributions of branch number and length

We next asked whether this local binary branching rule, inferred from the filament population, is sufficient to capture the variability observed in branching patterns among individual filaments. To answer this, we used a computational modeling approach based on the L-system formalism. L-systems allow to simulate the development of self-similar structures, such as compound branching structures, making them an ideal tool for modeling plant architecture^24–26^. In L-systems, a plant structure is represented as a string of modules, each corresponding to a distinct biological component. Lateral branches are encoded using brackets within the string, which specify the hierarchical dependencies between components. Development is simulated by iteratively applying production rules that dictate the fate of each module at every iteration step.

We thus defined a model of filament development (1) with two modules, A and C, representing apical and sub-apical cells, respectively. The simulation starts with a single apical cell A. At each iteration step, apical cells divide to generate a sub-apical cell (A *→* CA), and sub-apical cells produce a side-branch with probability *p* (C *→* C[A]). Since production rules are applied synchronously, a newly produced sub-apical cell cannot initiate a side-branch during the same iteration step. This constraint accounts for the observed lack of branching in the first sub-apical cell.

- **Modules** A, C
- **Axiom** A
- **Production rules**

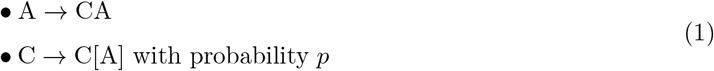

We implemented the local binary branching rule numerically using a branching probability *p* = 0.5 and simulated over 2000 virtual filaments (Figure 3A). Analyzing their branching patterns with the same metrics used for real filaments, we found that both side-branch number and length increased linearly along the filament axis with slopes of 0.5 and 0.6 (Figures 3B-3D), closely matching the experimentally measured values of 0.49 and 0.52, respectively (Figures 2B-2D). These results confirm that our probabilistic model accurately captures key features of filament branching observed at the population level. However, because this agreement reflects population averages, it may obscure variability among individual filaments. Consistent with this, differences remain between the simulated and biological datasets, as evidenced by comparing the corresponding boxplots in Figures 2B and 3B, and Figures 2C and 3C.

**Figure 3:**
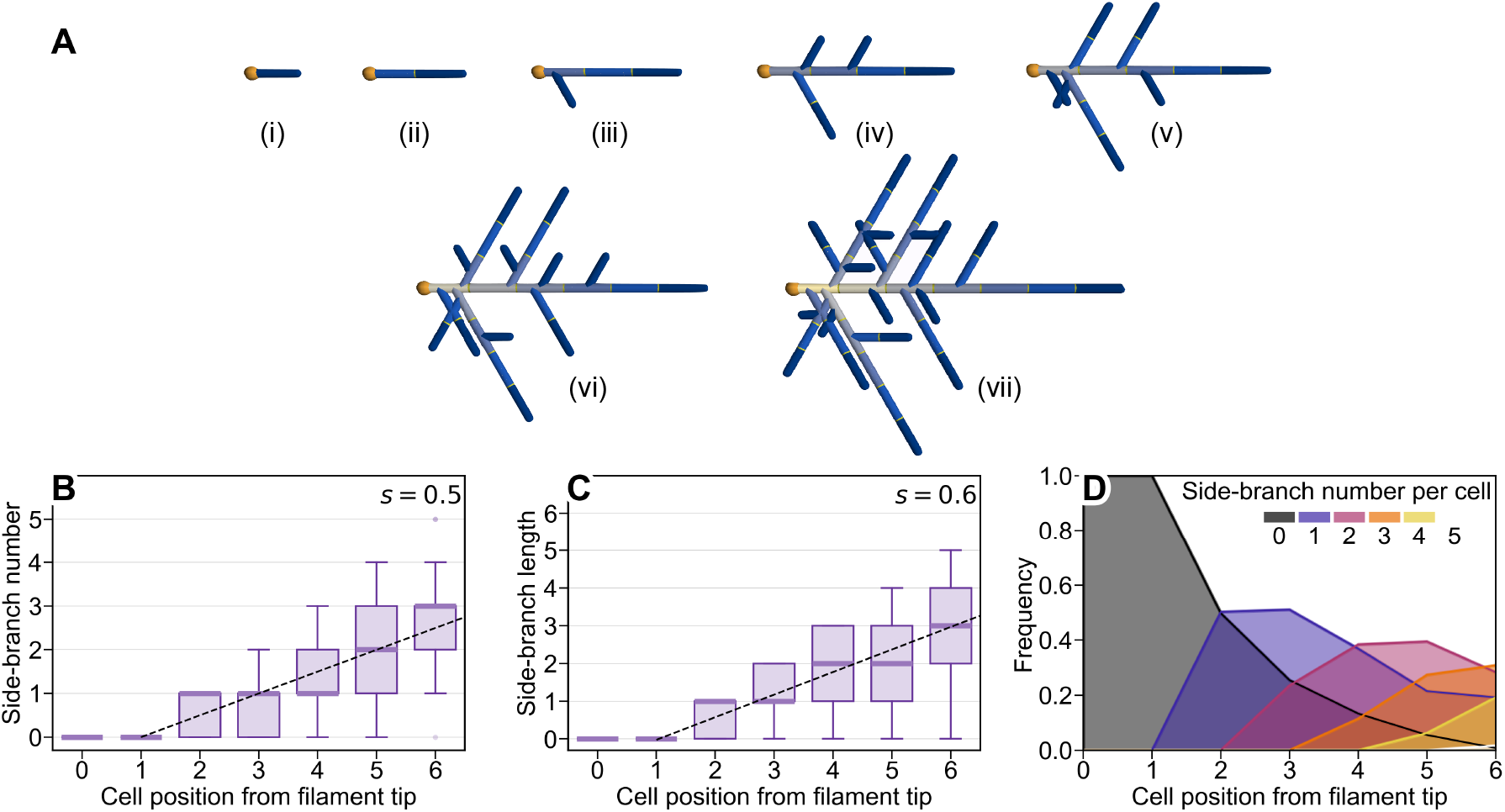
Side-branch length and number distribution can be replicated with a probabilistic model. **(A)** Example of a single simulation of the model described in Model (1), with branching probability *p* = 0.5, over 6 derivation steps. The color code represents cell position from the filament tip, from dark blue to yellow. Corresponding L-strings are (i) A, (ii) CA, (iii) C[A]CA, (iv) C[A][CA]C[A]CA, *etc*. **(B)** Box plots showing that the side-branch number per cell as a function of cell position from the tip in simulated filaments increased approximately linearly with slope *s* = 0.5. **(C)** Box plots showing that the distribution of side-branch length as a function of cell position from the tip in simulated filaments increased approximately linearly with slope *s* = 0.6, consistent with the observed data (*s* = 0.52, Figure 2C). **(D)** Intensity diagram of the side-branch number per cell as a function of cell position from the tip in simulated filaments, showing similar trends to the biological data (Figure 2D).

### Additional regulation shapes basitonic branching patterns in individual filaments

We hypothesized that these differences reflect deviations from strictly basitonic branching patterns (Figure 2A). To test this hypothesis and characterize the nature of these deviations, we quantified the cumulative side-branch length (CL, expressed in cell number) at each parent cell along individual filaments. To compare CL values across single filaments, we defined ΔCL as the difference between two consecutive CL values from the apex to the base, such that ΔCL_*i*_ = CL_*i*+1_ *−* CL_*i*_, where *i* corresponds to cell position from the filament tip (Figures 4A-B). ΔCL values were calculated starting from the first sub-apical cell with at least one side-branch. In principle, in a filament with strictly basitonic branching, ΔCL values are always positive, reflecting that younger parent cells have fewer side-branch cells than older ones (Figure 4A). Conversely, filaments with non-strictly basitonic branching show negative ΔCL values, indicating that older parent cells have fewer side-branch cells than younger ones (Figure 4B). In experimentally observed filaments, we calculated that 12% of ΔCL values were negative, indicating minor deviations from strictly basitonic branching patterns (Figure 4C). In contrast, 24% of ΔCL values were negative in simulated filaments, suggesting that the probabilistic model overestimates these deviations (Figure 4D). These results indicate that, although the probabilistic model captures much of the variability among individual filaments, an additional level of regulation is required to fully account for the greater branching regularity observed in real filaments compared with simulations.

**Figure 4:**
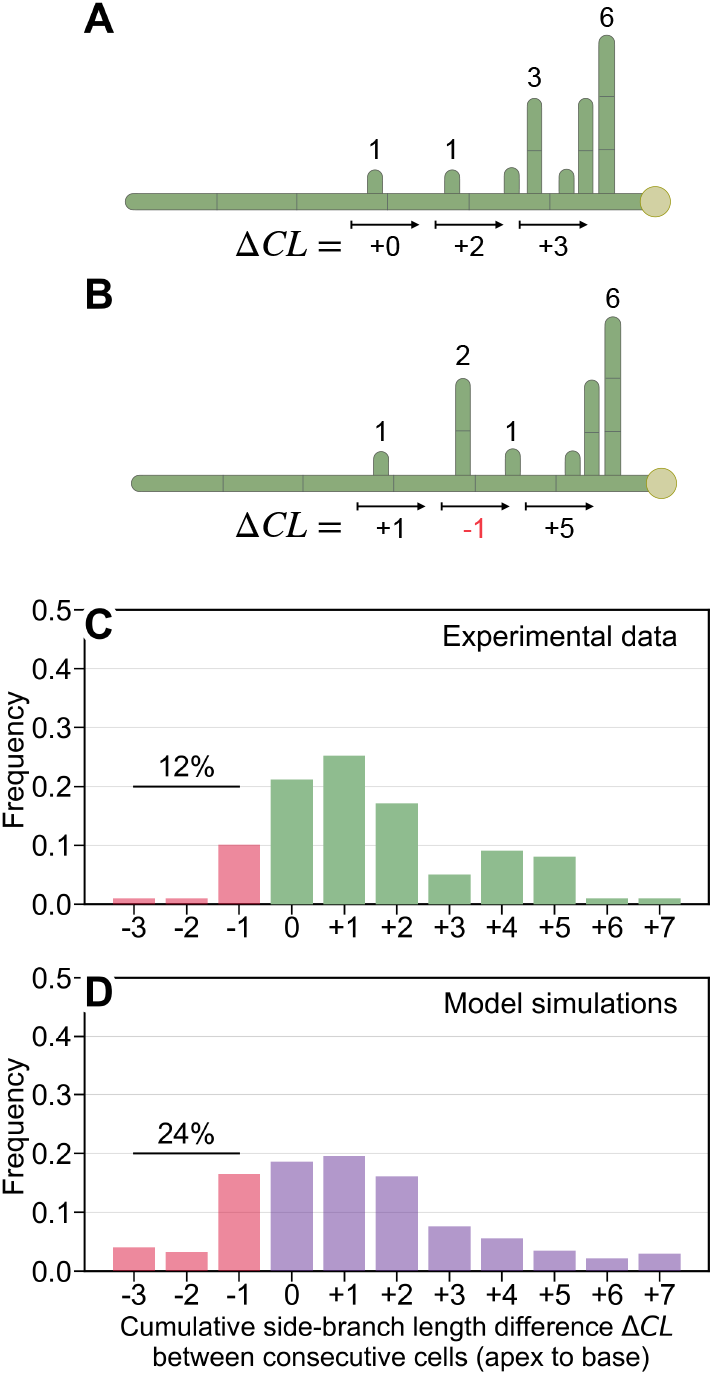
Branching patterns in individual filaments exhibit more regularity than expected under random branching. **(A)** Schematic diagram of a filament showing that the cumulative side-branch length differences (Δ*CL* values) are positive for basitonic branching patterns. **(B)** Schematic diagram of a single filament showing that deviations from strictly basitonic branching patterns are captured by negative Δ*CL* values (in red). **(C)** Histogram of frequencies of cumulative side-branch length differences between consecutive cells from apex to base (as represented in A-B), showing that sporeling filaments exhibit minor deviations from basitonic branching gradients, with 12% of negative differences. **(D)** Histogram of frequencies of cumulative side-branch length differences between consecutive cells from apex to base (as represented in A-B), showing that simulated filaments deviate more often from basitonic branching gradients than observed filaments, with 24% of negative Δ*CL* values.

## Discussion

This work aimed to identify developmental rules driving branching morphogenesis in the filamentous body of the moss *Physcomitrium patens*. We have taken a quantitative approach combining light-sheet microscopy, 3D image analysis and computational modeling, which revealed novel characteristics of branching patterns that could be recapitulated by a probabilistic model based on simple rules.

Consistent with previous reports investigating filament development at later stages, we found that filaments in early sporelings typically display a triangular morphology, with side-branch length increasing from apex to base. Side-branch number also increased along the apical-basal axis, reaching up to four side-branches per cell in basalmost positions. This observation indicates that filaments have a higher branching potential than previously thought^13,15^, possibly reflecting our experimental conditions. Notably, filaments were grown in liquid medium, whereas they are typically cultivated on solid medium in other studies^13,16^, suggesting that aquatic conditions may favour branch formation. Given that *P. patens* naturally grows in partially submerged environments^27^, this enhanced branching could reflect an adaptive physiological strategy promoting development under such conditions; however, this hypothesis remains to be tested. Additionally, our analysis focused on early developmental stages, whereas most previous studies have examined more mature gametophytes. The higher branch density observed may therefore reflect the onset of branching before inhibitory mechanisms are fully established. Consistent with this hypothesis, gradients of the phytohormone auxin – a major regulator of branching in plants – are progressively established during protonemal development^16,28^. Another notable feature was the near-absence of side-branches on the first sub-apical cell. While the underlying mechanism remains unknown, this feature may reflect a delayed acquisition of maturity and branching competence. Alternatively, it is also reminiscent of apical dominance, a well-described branch repression mechanism mediated by auxin in plant shoots^11,29,30^. The repeated deployment of apical dominance to regulate branch patterning across distantly related species and distinct life cycle phases underscores its evolutionary significance^15^. This observation motivates further quantitative investigation of this mechanism in moss filaments, particularly to clarify the relationship between local auxin levels and branch density.

Our study further demonstrates that branching patterns in early moss filaments are largely captured by a probabilistic model based on three simple rules: (1) apical cells divide periodically, (2) the first sub-apical cell cannot branch, and (3) sub-apical cells branch with probability *p* between successive apical cell divisions. The strength of this model lies in its simplicity, relying on a single parameter *p*, fitted here to 0.5, indicative of a highly entropic system. Beyond capturing the main features of our experimental data, this model provides a first quantitative framework for comparing filament branch patterning at the whole sporeling level across developmental time, genetic backgrounds and environmental conditions. More broadly, it contributes to establishing a unifying theoretical framework to compare macroscopic rules underlying branching across species, thereby advancing our understanding of the emergence of convergent biological forms. For instance, a provocative study combining quantitative imaging and modeling of branching morphogenesis in animal organs has proposed a “unifying theory of branching morphogenesis”^31^, whereby growing tips elongate and divide randomly, and existing branches inhibit nearby tip growth via diffusible compounds, resulting in a tight control of branch density. Similar density-dependent control of branching morphogenesis has been identified in arbuscular mycorrhizal fungi^32^, suggesting that such mechanisms may be shared across kingdoms. In *P. patens*, putative CCD8-derived strigolactones are secreted by filaments into their environment to inhibit their own branching and regulate sporeling expansion^33,34^. While our study suggests that cell-autonomous branching decisions underlie branch patterning in early sporelings, our modeling framework could be extended to explore whether density-dependent lateral inhibition contributes to filament patterning in *P. patens* and to assess the extent to which branching principles identified in animals and fungi extend to the plant kingdom^15^.

## Material and Methods

### Experimental model

All experiments were performed in the Gransden strain of *Physcomitrium patens* genetically transformed with the PpEF1*α*::acyl-YFP plasmid using PEG-mediated protoplast transfection^30^. To construct the PpEF1*α*::acyl-YFP plasmid, the acyl-YFP sequence^35^ was sub-cloned into the pDONR207 vector (ThermoFisher Scientific) via BP Gateway reaction and transferred to the pT1OG vector (https://www.nibb.ac.jp/evodevo/PHYSCOmanual/11.6_revised_170727.htm) via LR Gateway reaction, as described in (Lin *et al*., under revision)^36^.

### Tissue culture and growth conditions

Moss colonies were initiated from protonemal cultures and cultivated in petri dishes containing BCDAT medium supplemented with 7 g/L agar (Plant Agar, Ref P1001, Duchefa Biochemie, NL) and overlaid with cellophane disks. Spores were grown in liquid BCDAT medium. BCDAT culture medium was composed of 250 mg/L MgSO_4_.7H_2_O, 250 mg/L KH_2_PO_4_ (pH 6.5), 1010 mg/L KNO_3_, 12.5 mg/L, FeSO_4_.7H_2_O, 920 mg/L di-ammonium tartrate (C_4_H_12_N_2_O_6_), and 0.001% Trace Element Solution (0.614 mg/L H_3_BO_3_, 0.055 mg/L AlK(SO_4_)_2_.12H_2_O, 0.055 mg/L CuSO_4_.5H_2_O, 0.028 mg/L KBr, 0.028 mg/L LiCl, 0.389 mg/L MnCl_2_.4H_2_O, 0.055 mg/L CoCl_2_.6H_2_O, 0.055 mg/L ZnSO_4_.7H_2_O, 0.028 mg/L KI and 0.028 mg/L SnCl_2_.2H_2_O), and supplemented with CaCl_2_ added to a 1 mM concentration after autoclaving. Standard growth conditions were 23°C under a 16h light/8h dark cycle, at 50-150 *µ*mol m^*−*2^s^*−*1^. The liquid medium was continuously agitated on an orbital shaker at 50 rpm. Spores were imaged after 4-7 days of culture.

### Sporophyte induction

To obtain spores, moss colonies were transplanted onto sterile peat pellets and cultivated for one month at 23°C under long days conditions (16 hours light/8 hours dark). Gametangia formation was then induced by transfering moss cultures to low temperature (16°C) and short day conditions (8 hours light/16 hours dark). Sexual reproduction was induced by submerging the peat pellets in sterile MilliQ-H_2_O. Once sporophytes were fully developed, individual capsules were isolated and sterilized by immersion in 2.5% sodium hypochlorite for 5 minutes, followed by three rinses with sterile MilliQ-H_2_O. A single capsule was disrupted and spores were diluted in 15mL MilliQ-H_2_O. The resulting spore solution was stored at 4°C until use.

### Light-sheet imaging

Spore germinations (or sporelings) were mounted in glass capillaries (1 mm inner diameter) in 2% low-melting-point agarose. The sample chamber was filled with MilliQ-H_2_O. Imaging was performed with the Zeiss Lightsheet Z1 microscope using a 20x detection objective (W Plan-Apochromat 20x/1.0 Corr DIC M27 75 mm) and two 10x/0.2 illumination objectives. To reduce shadow artifacts, laser pivoting was enabled. Light sheet position was adjusted at the beginning of each acquisition batch. Light sheet thickness was set to the optimal value, typically 4.55 *µ*m. YFP excitation was performed using a 488 nm laser at 20% power with 30 ms exposure, and chlorophyll excitation was performed using the same laser at 0.8% power with 20 ms exposure. Channel alignment was performed directly after image acquisition using the ZEN blue software (Zeiss, Germany).

### Image segmentation

For image segmentation, an automatic pipeline presented in Mosaliganti *et al*., 2012^37^ and implemented in Arivis Vision4D 3.6 was used. This pipeline consisted of a pre-processing step referred to as “Membrane detection”, and a segmentation step referred to as “Membrane-based segmentation”. The membrane detection algorithm relied on tensor voting to detect planar signal and improve the image to enhance the contrast and integrity of the membrane signal. This algorithm was parameterized with the size of the neighborhood used to estimate signal planarity at each voxel and the maximum size of signal gaps to be filled. Neighborhood size was set to 1-1.5*µ*m, corresponding to the average membrane width, and maximum gap size was set to 0.5*µ*m, corresponding to approximately two voxels. Segmentation was parameterized with a minimum signal intensity threshold used to binarize the image between membrane and background and a split sensitivity affecting the ratio of over and under-segmentation. The minimum signal intensity was set to 20 and the split sensitivity was set to 25-35%. Objects smaller than 1000*µ*m^3^ were removed from the segmentation results. Using the Arivis Vision4D viewer and associated curation tools, and based on careful inspection of the original intensity images, labelled images were corrected to merge over-segmented cells, dissociate under-segmented cells, and manually draw missing cells or cell portions. Original datasets are available upon request to the corresponding author.

### Tree representation

To reconstruct the cellular tree of sporelings, with nodes corresponding to cells and edges corresponding to cell divisions, the cell adjacency graph was extracted from the 3D labelled images using `timagetk`, a Python-implemented image analysis library (https://mosaic.gitlabpages.inria.fr/timagetk/). The cell adjacency graph was manually corrected to delete edges that did not correspond to actual cell divisions, until the tree structure of cell organization was recovered. The spore was identified from the bright field channel of the sporeling images and was set as the root of the tree. Tree implementation was done using `treex`, a Python-implemented library dedicated to manipulating rooted trees^38^ (https://mosaic.gitlabpages.inria.fr/treex/).

### Branching order estimation

Estimating branching order in sporelings required identifying the main axis and side-branches among the children of each node. This was done manually based on visual cues such as the overall curvature of filaments and the orientation of cell divisions in the intensity images. In 46 out of 193 cases, the main axis could not be confidently identified from visual cues and was set to correspond to the cell at the root of the largest subtree.

### Chloroplast density estimation

Chloroplasts were segmented using the “Blob finder” algorithm implemented in Arivis Vision4D 3.6, applied on the channel image corresponding to chlorophyll autofluorescence. This algorithm was parameterized with the average chloroplast diameter, estimated to 3.7*µ*m, a probability threshold expressed in percent of the intensity range of the complete object probability map, set to 20%, and a split sensitivity set to 65%. Based on the identification of cells obtained with the membrane-based segmentation, segmented chloroplasts were assigned to each cell. Chloroplast density was defined for each cell as the total chloroplast volume over the cell volume.

### Cell length estimation

For each segmented cell, its skeleton was first determined using the Pythonimplemented function `skimage.morphology.skeletonize`, which implements the algorithm presented in Lee *et al*., 1994^39^. The longest skeleton path inside that cell was defined as the cell length.

### Model implementation and simulation

The probabilistic model of filament branching was numerically implemented using L-Py^26^, and simulation results were analyzed using the `treex` library^38^. A set of 2180 simulated filaments was analyzed. The number of iteration steps per simulation was set to match the primary filament size proportions between observed and simulated filaments. Code is available upon request to the corresponding author.

## Acknowledgments

This work was financially supported by the CNRS (ATIP-Avenir grant to Y.C., Ph.D. fellowship to J.A.-S.). We acknowledge the contribution of SFR Santé Lyon-Est (UAR3453 CNRS, US7 Inserm, UCBL) facility CIQLE (a LyMIC member) and Denis Ressnikoff. J.A.-S. acknowledges the contribution of Laure Mancini for her assistance with sporophyte induction, Jonathan Legrand and Guillaume Cerutti for their assistance with the `timagetk` image analysis Python library, and Romain Azais for his assistance with the `treex` Python library.

## Author contributions

Conceptualization: J.A.-S., C.G., and Y.C.; Investigation: J.A.-S., with support from B.C. for light-sheet imaging; Software: J.A.-S.; Formal Analysis: J.A.-S.; Writing – Original Draft: J.A.-S. and Y.C.; Writing – Review and Editing: J.A.-S., C.G., and Y.C.; Supervision: C.G. and Y.C.; Funding Acquisition: Y.C.

## Conflicts of interest statement

The authors declare no conflicts of interest.

## Declaration of AI assistant use

The authors, who are non-native English speakers^40^, used an AI assistant to enhance the readability of the English text of the manuscript. The content was carefully reviewed by the authors who take full responsibility for the integrity of the publication.

## Notes

### Competing Interest Statement

The authors have declared no competing interest.

