## Supplemental information for "Near-equiprobable binary branching decisions underlie filament patterning in the moss *Physcomitrium patens*"

[1]Laboratoire Reproduction et Développement des Plantes, CNRS, ENS de Lyon, Université Claude Bernard Lyon 1, INRAE, INRIA, Lyon 69007, France
[2]Centre d'Imagerie Quantitative Lyon-Est, SFR Santé Lyon-Est (UAR3453 CNRS, US7 Inserm, UCBL), Lyon 69007, France
[3]These authors contributed equally
[4]Corresponding author

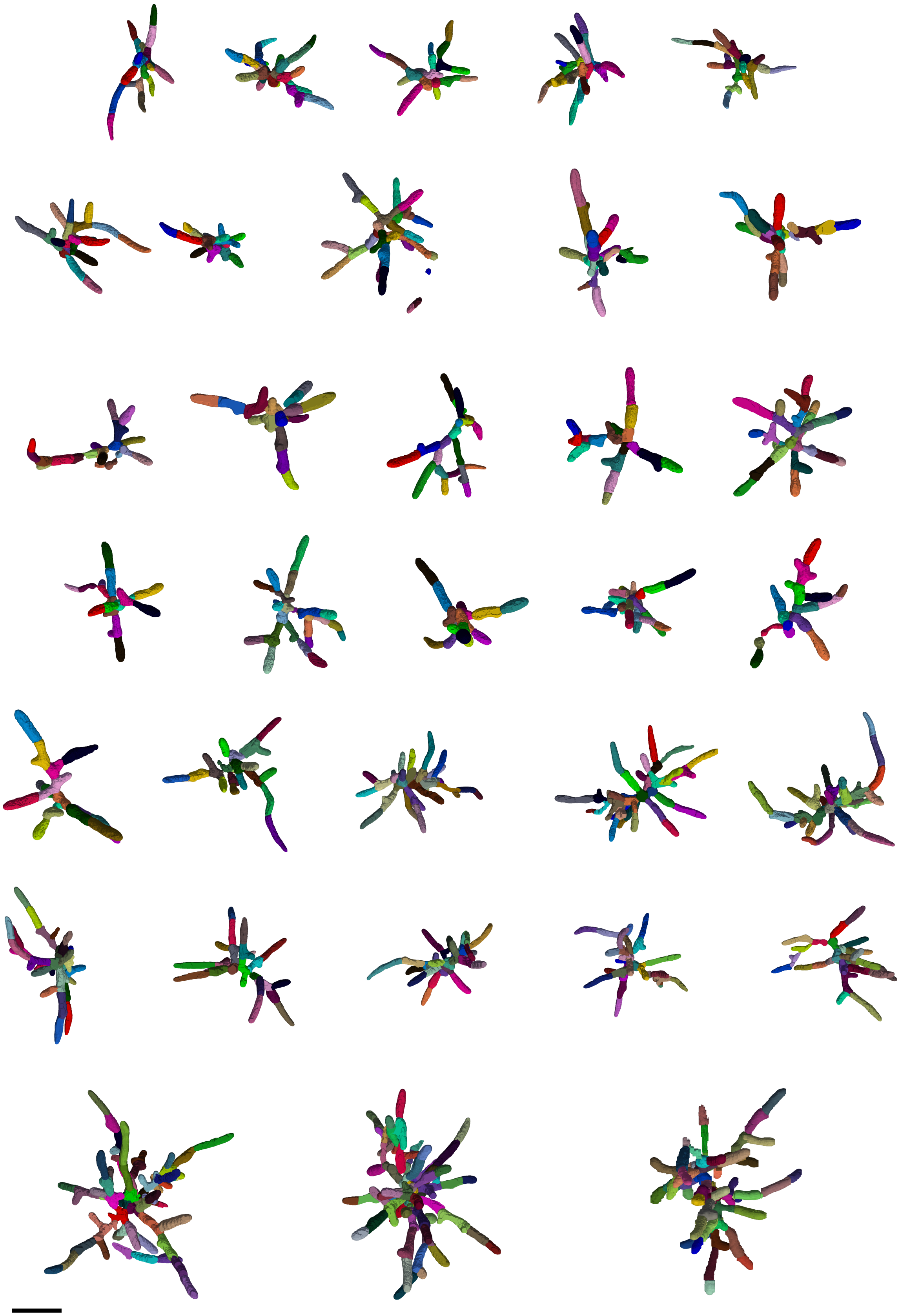

**Figure S1: Filament segmentation overview.** Overview of cell segmentation results of all sporelings included in the dataset analyzed in this study. Scale bar = $100 \mu m$.

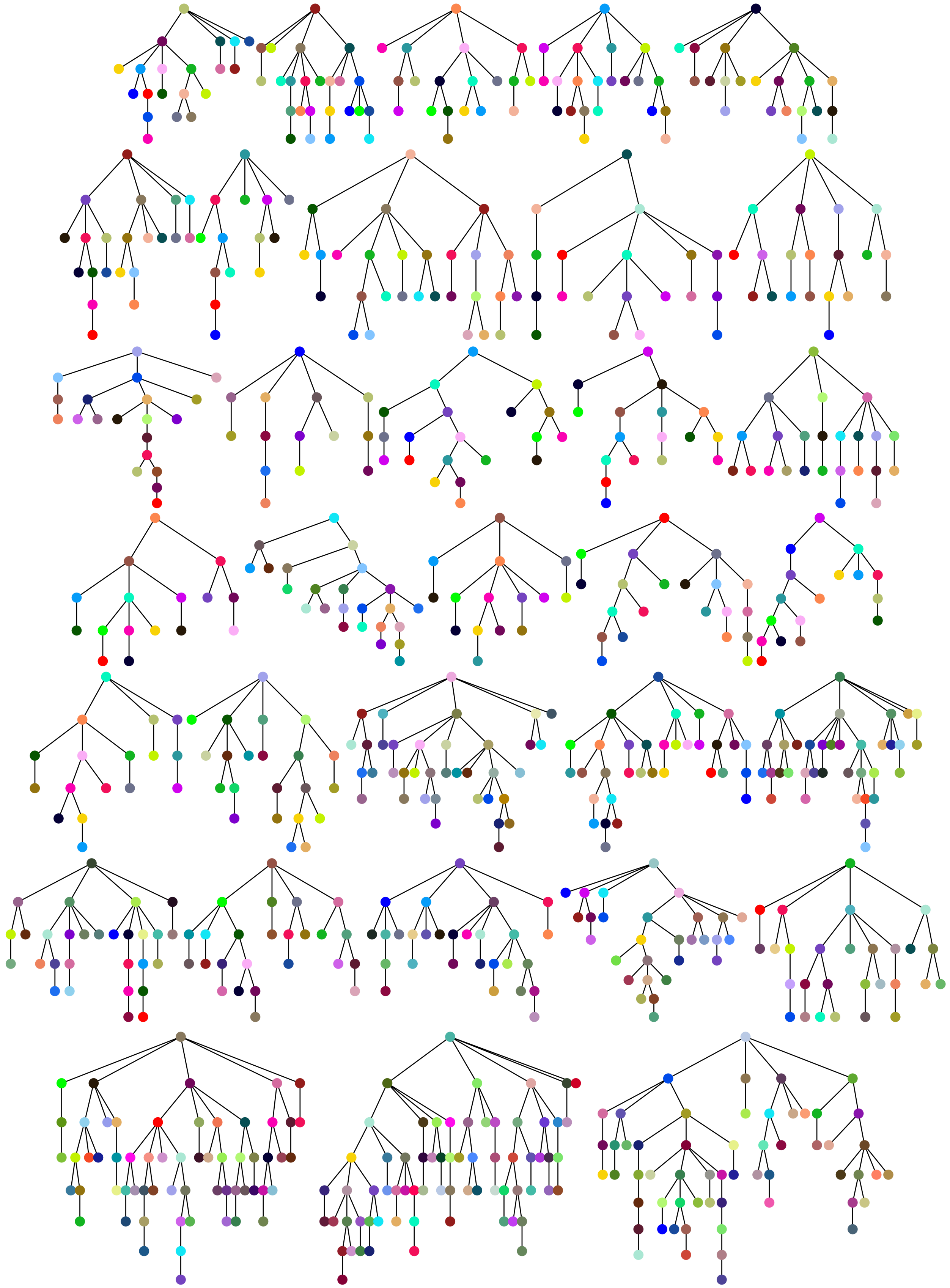

**Figure S2: Overview of tree-like structures representing the architecture of analyzed sporelings.** *(Legend next page.)*

*(Figure on previous page.)* Overview of the tree-like structure of all sporelings constituting the dataset analyzed in this study. Nodes represent cells and edges represent cell divisions. Trees are rooted at the spore, which is represented as the upmost node. Note that branching order is not represented and beyond root upmost position, spatial organization of nodes within each tree is not informative. Trees are spatially organized consistently with Figure S1. Node colors correspond to cell labels consistently with Figure S1.

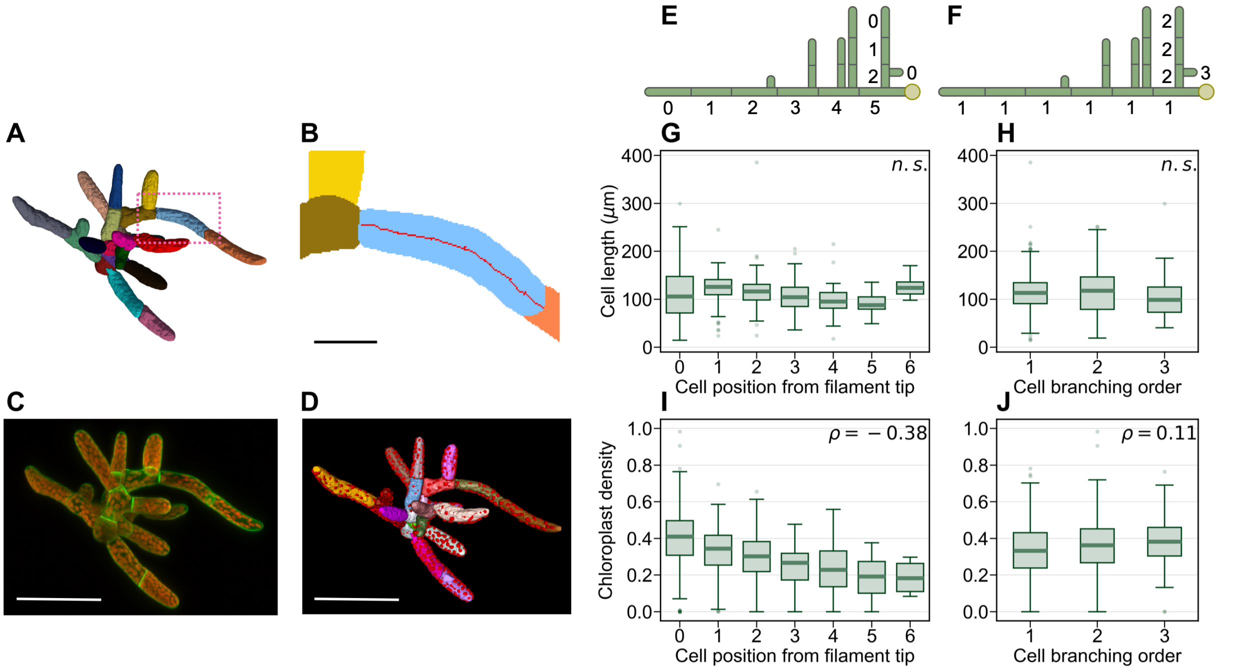


**Figure S3: All sporelings were imaged before the caulonemal transition. (A)** Segmentation of a single sporeling. **(B)** 2D projection of the skeletonization result of the blue cell framed in (A). 3D skeletonization results were used to estimate cell length in filaments. **(C)** Maximum intensity projection of a light-sheet micrograph of a sporeling. Membrane signal intensity is shown in green and chloroplast autofluorescence in red. **(D)** Segmentation of individual chloroplasts (in red) of the sporeling shown in (C), overlaid on segmented cells. Based on segmentation results, chloroplast density was estimated for each cell as the cumulated volume of chloroplasts divided by the volume of the corresponding cell. **(E-F)** Schematic representation of a sporeling filament, with cells numbered according to distance from the filament tip (E) or cell branching order (F). **(G-H)** Boxplots of cell length distribution as a function of cell position from the filament tip (G) and of cell branching order (H), showing that no cell length gradient was detected in the analyzed sporelings. **(I-J)** Boxplots of chloroplast density distribution as a function of cell position from the filament tip (I) and of cell branching order (J), showing that chloroplast density negatively correlated with distance from the tip and positively correlated with cell branching order. Pearson correlation coefficients $\rho$ are displayed if statistically significant ($p < 0.05$). Scale bars = 20 $\mu$m (B), 100 $\mu$m (C,D).
